# Pod and seed trait QTL identification to assist breeding for peanut market preferences

**DOI:** 10.1101/738914

**Authors:** Carolina Chavarro, Ye Chu, Corley Holbrook, Thomas Isleib, David Bertioli, Ran Hovav, Christopher Butts, Marshall Lamb, Ronald Sorensen, Scott A. Jackson, Peggy Ozias-Akins

## Abstract

Although seed and pod traits are important for peanut breeding, little is known about the inheritance of these traits. A recombinant inbred line (RIL) population of 156 lines from a cross of Tifrunner x NC 3033 was genotyped with the Axiom_Arachis1 SNP array and SSRs to generate a genetic map composed of 1524 markers in 29 linkage groups (LG). The genetic positions of markers were compared with their physical positions on the peanut genome to confirm the validity of the linkage map and explorethe distribution of recombination and potential chromosomal rearrangements. These traits were phenotyped over three consecutive years for the purpose of developing trait-associated markers for breeding. Forty-nine QTL were identified in 14 LG for seed size index, kernel percentage, seed weight, pod weight, single-kernel, double-kernel, pod area and pod density. Twenty QTL demonstrated phenotypic variance explained (PVE) greater than 10% and eight more than 20%. Of note, seven of the eight major QTL for pod area, pod weight and seed weight (PVE >20% variance) were attributed to NC 3033 and located in a single linkage group, LG B06_1. In contrast, the most consistent QTL for kernel percentage were located on A07/B07 and derived from Tifrunner.

## INTRODUCTION

Peanut (*Arachis hypogaea* L.), also referred to as groundnut, is an important legume for human nutrition due to its high levels of protein and oil. It is one of the most important crop legumes in the world with an annual production of 42.9 million metric tons in 2016 (FAO 2017). Seed size and quality are important for breeding and production, thus, a more mechanistic understanding of pod development and seed maturation would benefit the improvement of these traits. During pod development, seed filling plays an important role due to the translocation of organic and inorganic compounds and is an important yield component (Shiraiwa *et al*. 2004; Madani *et al*. 2010; El-Zeadani *et al*. 2014). During seed maturation, the pod filling process is complete when the seeds accumulate nutrients and reach their maximum volume (Mahon and Hobbs 1983; Habekotté 1993; Imsande and Schmidt 1998; Clements *et al*. 2002).

Cultivated peanut is an allotetraploid (2n = 4x = 40) with a genome size of 2.7 Gb, approximately the sum of the two diploid A- and B-genome progenitors, *A. duranensis* and *A. ipaensis*, respectively (Samoluk *et al*. 2015). Cultivated peanut was derived from the hybridization of these two diploids (Kochert *et al*. 1996; Fávero *et al*. 2006; Seijo *et al*. 2007; Robledo *et al*. 2009; Robledo and Seijo 2010; Moretzsohn *et al*. 2013; Bertioli *et al*. 2016) that diverged from each other ∼2.2 – 3.5 million years ago (Nielen *et al*. 2012; Moretzsohn *et al*. 2013; Bertioli *et al*. 2016). The polyploidization event was very recent, at most ∼9-10 thousand years ago (Bertioli *et al*. 2016) which reproductively isolated cultivated peanut from its wild diploid relatives.

This evolutionary history has resulted in low levels of genetic variation (Kochert *et al*. 1991) within tetraploid peanut and high collinearity between the A and B sub-genomes (Moretzsohn *et al*. 2009; Guo *et al*. 2012; Shirasawa *et al*. 2013; Bertioli *et al*. 2016, 2019); thus, gene discovery for breeding is challenging (Stalker and Mozingo 2001; Holbrook *et al*. 2011; Chu *et al*. 2016; Guo *et al*. 2016). Furthermore, the low polymorphism rates and similarity between the two subgenomes of cultivated peanut delayed the development and implementation of genotyping tools and the identification of markers for breeding (Holbrook *et al*. 2011; Shirasawa *et al*. 2012; Koilkonda *et al*. 2012; Clevenger *et al*. 2017). To avoid the challenges of polyploidy and low levels of polymorphism in cultivated peanut, a few medium density genetic maps of diploid relatives have been constructed (Nagy *et al*. 2012; Bertioli *et al*. 2014; Leal-Bertioli *et al*. 2015) including consensus maps for the A and B genomes based on wild species (Shirasawa *et al*. 2013).

In the past few years, however, genome sequences for peanut (Bertioli *et al*. 2019) and its progenitors (Bertioli *et al*. 2016) along with advances in the SNP identification and detection (Clevenger and Ozias-Akins 2015) have resulted in thousands of SNP markers (Pandey *et al*. 2017; Clevenger *et al*. 2017, 2018). Mapping with SNP markers has led to more saturated maps in cultivated peanut with the number of mapped loci ranging from 772 SNPs to 8,869 SNPs (Zhou *et al*. 2014; Huang *et al*. 2016; Liang *et al*. 2017; Agarwal *et al*. 2018; Wang *et al*. 2018a, 2018b; Liu *et al*. 2019) and QTL identified reviewed by Ozias-Akins et al. (2017).

Mapping of seed and pod traits in bi-parental populations has included QTL analyses for pod and seed length, width, weight and number of seed per pod (Gomez Selvaraj *et al*. 2009; Fonceka *et al*. 2012; Shirasawa *et al*. 2012; Wu *et al*. 2014; Huang *et al*. 2015; Chen *et al*. 2016a, 2017, 2019; Luo *et al*. 2017, 2018; Wang *et al*. 2019, 2018a) as well as associations for pod and seed weight, number of seeds and pods per plant (Gomez Selvaraj *et al*. 2009; Ravi *et al*. 2010; Fonceka *et al*. 2012; Shirasawa *et al*. 2012; Wang *et al*. 2018a, 2019; Chen *et al*. 2019), shelling percentage (Faye *et al*. 2015; Huang *et al*. 2015; Chen *et al*. 2016a), pod maturity (Liang *et al*. 2009b; Gomez Selvaraj *et al*. 2009; Fonceka *et al*. 2012; Faye *et al*. 2015), and morphological traits such as pod constriction, thickness or seed coat color (Fonceka *et al*. 2012; Shirasawa *et al*. 2012). However, none of these studies included pod density as an indicator of pod filling on the yield components, and most of these studies were limited by the small number of markers (∼220-820 markers) (Liang *et al*. 2009a; Gomez Selvaraj *et al*. 2009; Ravi *et al*. 2010; Fonceka *et al*. 2012; Huang *et al*. 2015; Chen *et al*. 2016a, 2017; Luo *et al*. 2017, 2018; Wang *et al*. 2019), except Shirasawa et al. (2012) that included 1114 SSRs and, more recently, Wang et al. (2018a) which included 3630 SNPs.

In this study, a saturated genetic map was constructed using a set of recombinant inbred lines (RILs) from a cross of two peanut genotypes, Tifrunner x NC 3033. This population was phenotyped for seed and pod traits for three consecutive years. While seed and pod trait QTL have been identified in previous studies, none are associated with pod filling as a yield component. The hypothesis of this study states that the measurement of seed and pod traits such as kernel percentage and pod density as a measure of pod filling along with other traits such as individual pod and seed weight, number of seeds per pod and 16/64 percentage, a standard measure of the kernel size for commercial purposes (USDA 1997), will help us to identify novel QTLs and confirm previous QTLs found by other researchers. As a result, a linkage map including 1524 markers was constructed and forty-nine QTL were discovered for seed and pod traits, including eight major QTL. These results will enhance our ability to improve peanut seed quality and yield through molecular breeding by providing molecular markers for marker assisted selection (MAS).

## MATERIALS AND METHODS

### Plant material

A set of RILs derived from a cross of Tifrunner x NC 3033 was developed and roughly half were advanced in Tifton, Georgia and the remainder in Raleigh, North Carolina (Holbrook *et al*. 2013). NC 3033 (*Arachis hypogaea* L. subsp. *hypogaea* var. *hypogaea*) (Beute *et al*. 1976; Hammons *et al*. 1981) is a small-seeded Virginia type germplasm line with incomplete pod filling, while Tifrunner (*Arachis hypogaea* L. subsp. *hypogaea* var. *hypogaea*), a released cultivar, has more complete pod filling. NC 3033 is resistant to several diseases including stem rot (*Sclerotium rolfsii* Sacc.) and is one of the most cylindrocladium black rot (CBR) resistant genotypes identified (Hadley *et al*. 1979). However, NC 3033 has low seed grades and low % meat as compared to Tifrunner, an elite runner type characterized by large seeds and good grade (Holbrook and Culbreath 2007) (Fig. 1).

**Figure 1.**
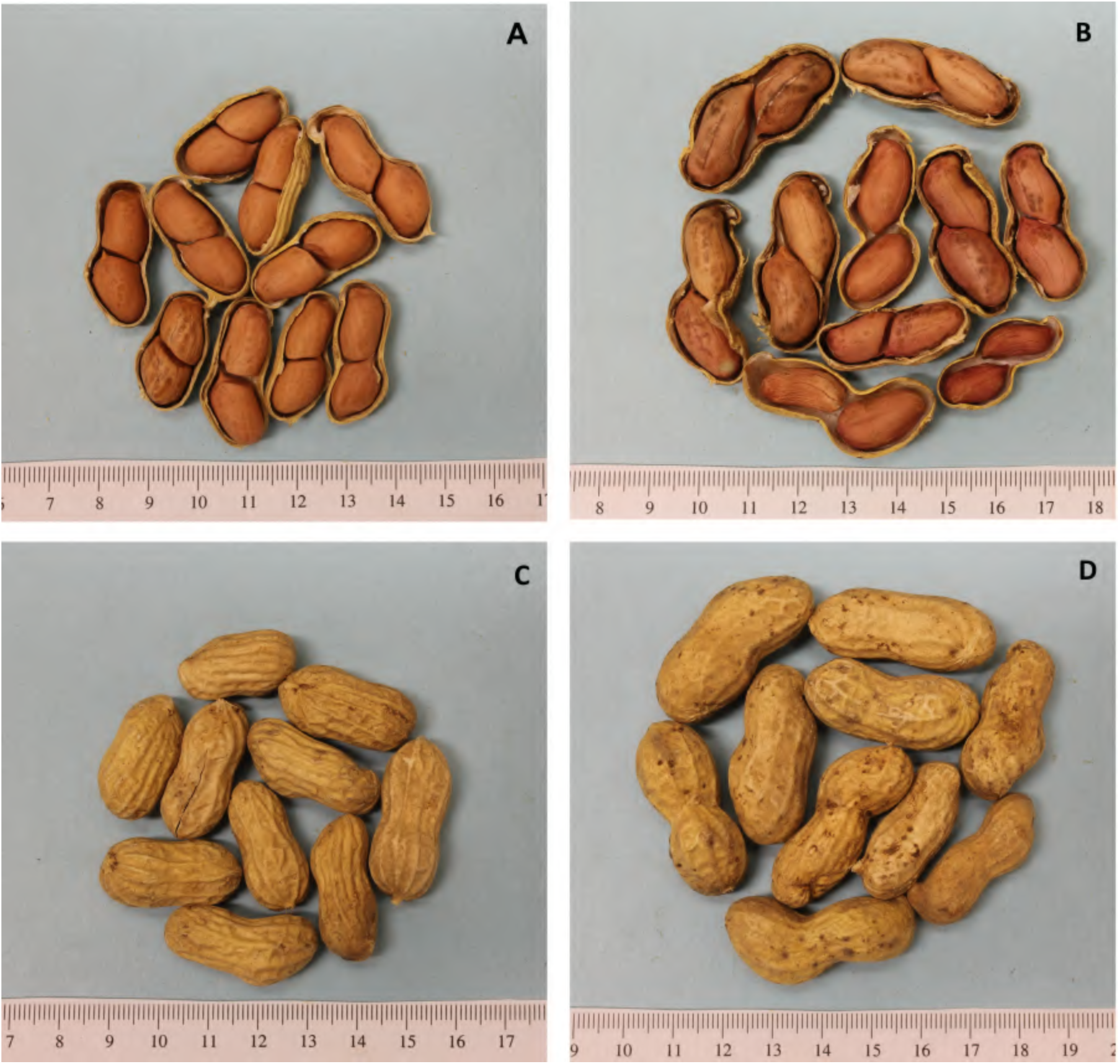
Seeds and pods from Tifrunner and NC 3033. **A, C.** Tifrunner, a commercial runner type in seed and pod size showing complete pod-filling. Note the proximity from the seeds to the border of the pods, which is a desirable commercial trait. **B, D.** NC 3033, a small seeded Virginia type showing incomplete pod-filling. Note how the seeds are loose and do not reach the border of the pods.

### Phenotyping of seed and pod traits

The Tifton-derived portion of the RIL population was planted for three consecutive years in Tifton, GA (USA) and phenotyping was conducted for 134 F_6:8_ RILs in 2013, 152 F_6:9_ RILs in 2014 and 160 F_6:10_ RILs in 2015 using a randomized complete block experimental design with three replicates and a plot size of 1.5 m x 1.8 m. For all years, 16/64 percentage (16/64P) as seed size index and kernel percentage (KP), also known as shelling percentage, were obtained using a BestRay X-ray grading machine. 16/64P is the percentage by weight of seeds that fall through a 16/64 x ¾ in screen retaining seeds with size of interest. KP and 16/64P are calculated as proportion of the sum of kernel weight and hull weight for 100 pods.

In 2014 and 2015, a subset (250 g of pods) was selected for each RIL to determine the variation in pod filling through phenotyping of individual pods. The pods were dried to approximately 10% moisture and then classified and counted based on the number of seeds per pod in single- (SP), double- (DP) and triple-kernel pods (TP). Due to a low number of triple-kernel pods found in only a few individuals, this trait was not used for the QTL analysis since the data was not transformable to follow a normal distribution. Subsequently, 10 randomly-selected double-kernel pods per line and replicate were shelled and the maturity was judged by the internal pericarp color (IPC) (Gilman and Smith 1977). The weight of the entire pod including the shell and the kernels was recorded (PW) and the weight of the two kernels was recorded (SdW) using the LabX Balance Direct 3.2 software and a digital scale. Ten half pods per line per replicate were scanned on both sides and analyzed using ImageJ (Rasband 2011) to determine pod area (PA) as a surrogate for pod volume according to Wu et al. (2015). The pod density (PD) (pod density = pod weight / pod area mm^2^) was calculated for the samples as a measure of pod filling.

### Statistical analysis

For all the phenotypic traits, Shapiro-Wilk and Anderson-Darling tests for normality of distribution were performed. When the data did not fit a normal distribution, outliers were removed and the data were transformed (e.g. logarithmic, square root or reciprocal values). Correlation coefficients between all the traits across years for the parents were calculated using Minitab 17 (Minitab® 17 Statistical Software 2010). Histograms, boxplots and analysis of variance for all the traits and years were plotted using R. Two-way ANOVAs for all the traits were made following the linear model method in R to identify significant differences between RILs, blocks and the interaction between RILs x years. Following the same model, broad sense heritability was determined by calculating (SS RIL) / (SS model – SS block), where SS corresponds to the sum of squares. To diminish the block effect for the analysis of variance, the year effect was calculated in a separate model including only year effects.

### Genotyping and map construction

The parents, Tifrunner and NC 3033, were included in a panel of genotypes sequenced by whole genome re-sequencing to identify the SNPs for the Affymetrix Axiom_Arachis SNP array containing 58,233 SNPs (Pandey *et al*. 2017; Clevenger *et al*. 2017). DNA of the parents and a set of 165 F_6:7_ RILs of the population planted in Tifton was extracted using the Qiagen DNeasy Plant mini kit® and sent to Affymetrix for genotyping. SNP calls were curated using the Axiom Analysis Suite Software® (Thermo Fisher Scientific Inc. 2016) based on the clustering of data for the entire population and the parents. Also included were 111 fluorescence tagged SSRs (Guo *et al*. 2012), previously used to genotype this population.

All RILs were checked for segregation distortion using a χ^2^ test and an expected 1:1 segregation ratio. Markers and RILs with more than 10% missing data were removed as well as the RILs with more than 20% heterozygote calls. A genetic map was constructed using JoinMap v4.1 (Van Ooijen 2006) with a minimum LOD of 3.0 and the Kosambi function. A graphical representation of the map was constructed using Mapchart v2.3 (Voorrips *et al*. 2002).

Linkage groups were identified and named based on the pseudomolecules of the tetraploid *A. hypogaea* genome cv. Tifrunner (Bertioli et al 2019; http://peanutbase.org). Marker locations were compared to SNP sequence positions on the pseudomolecules of the two ancestral diploid genomes (Bertioli *et al*. 2016; Clevenger *et al*. 2017). Confirmation of the loci positions was done manually and by BLASTN (*e* value < 1 × 10^−10^) of the SNP flanking sequences to the tetraploid reference genome, using an identity greater than 90%, alignment greater than 80% and fewer than three mismatches.

### QTL analysis

The normalized and average values from the three replicates of the phenotypic traits per year were used for QTL identification (File S1). Composite Interval Mapping was performed using WinQTL Cartographer v2.5_011 (Wang *et al*. 2012). The statistical significance of the QTL effects was determined using 1000 permutations with a 0.05 significance level. A graphical version of the map with QTL was constructed using Mapchart v2.3 (Voorrips *et al*. 2002). Naming of QTL follows the nomenclature of “q” as QTL, followed by the abbreviation of the trait, the last two digits of the year and the consecutive number of the QTL for that specific trait. The markers flanking the QTL were used to obtain the physical position from the *A. hypogaea* genome.

QTL were compared with previously reported QTL for seed and pod traits, based on physical and genetic locations. The flanking sequences of the markers linked to QTL or the fragment sequence of the QTL regions related to seed, pod and yield traits reported by Gomez Selvaraj et al. (2009), Fonceka et al.(2012), Chen et al. (2016a, 2019), Luo et al. (2018) and Wang et al. (2018a) were extracted from the two diploid progenitors. BLASTN was performed with e-value 1e −10, gap open 5, gap extend 2, penalty −2, against the *A. hypogaea* genome sequence. The first hit was taken for comparison of LG and position. The position of the hit was compared with the position of the QTL reported in this study to determine possible overlap. In addition, comparisons were made with the integrated QTL described by Chen et al. (2017), based on the reported physical position on the diploid genome progenitors and compared to the physical position of the QTL in this study, also based on the diploid genomes following the same BLASTN parameters.

### Data availability

The phenotypic information, the linkage map information and the genotyping used for map construction are described in Supporting Information, File S1. The phenotypic information includes the measurement and transformation method. The linkage map information and genotyping include the genetic and physical positions of the markers plus the GenBank accession ID for the SSRs available. Table S1 describes the previous QTL identified in cultivated peanut used in this study for comparison.

## RESULTS

### Seed and pod phenotypes in the RIL population

NC 3033, although a small-seeded Virginia type peanut with incomplete pod filling (e.g. R7 stage (Boote 1982) in Fig. 1), has larger seeds than Tifrunner. Phenotypic data of the parents and the RIL population were collected over three years using a randomized complete block design (Table 1). We observed a large block effect in 2015 that can be attributed to moisture (rain) after harvest where two replicates (2 and 3) were infested with mold that affected pod weight and density (Fig. 2). For most of the phenotypic data, we were able to obtain normal distributions (Fig. 3).

**Figure 2.**
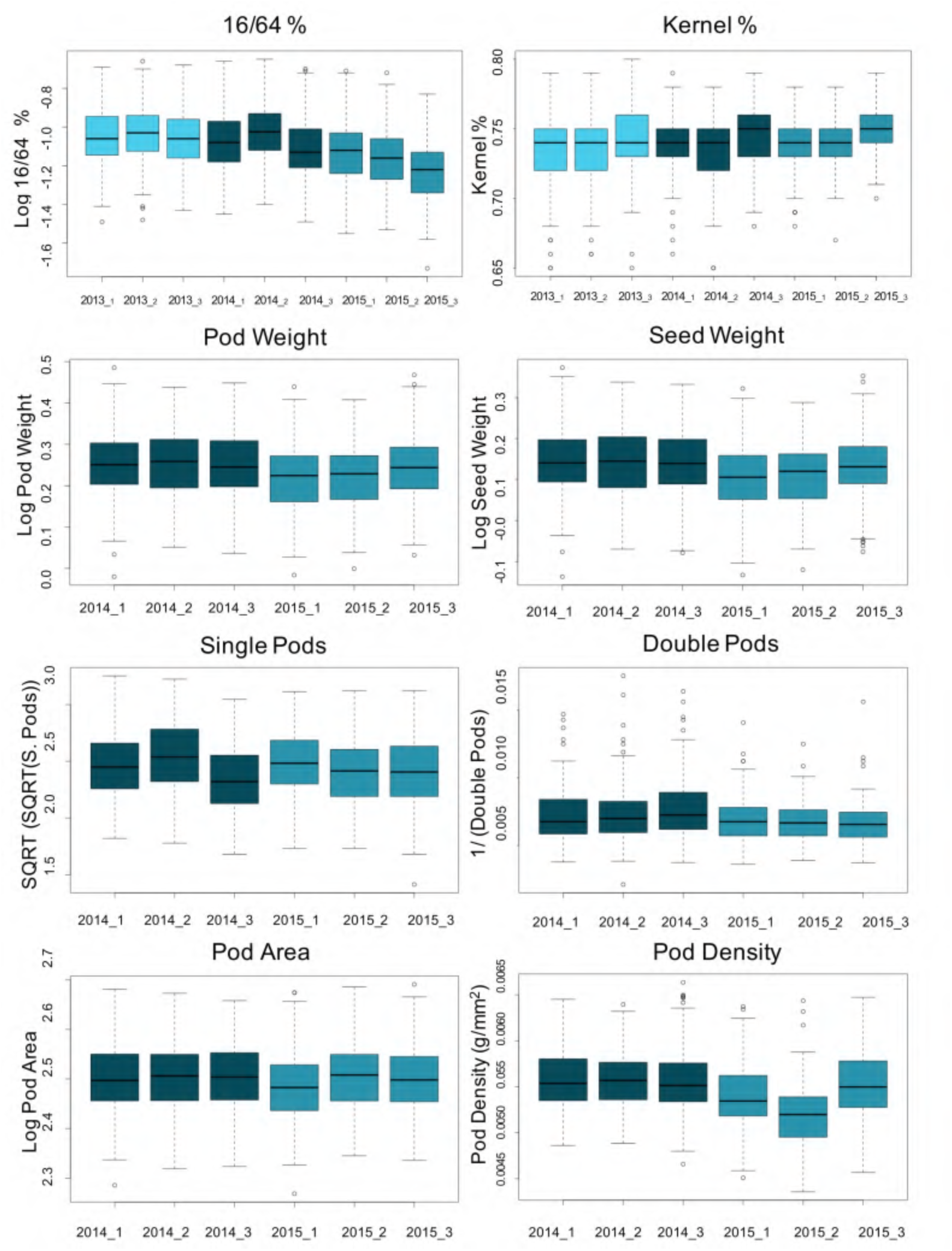
Boxplots for seed and pod traits across years, blocks and replicates within years, based on the normalized data. y-axis indicates the original metric or the normal-transformed of the trait value and the x-axis the years and replicates within a year. The color of the boxes indicates different years. Light sky blue, 2013; dark blue, 2014, teal blue, 2015.

**Figure 3.**
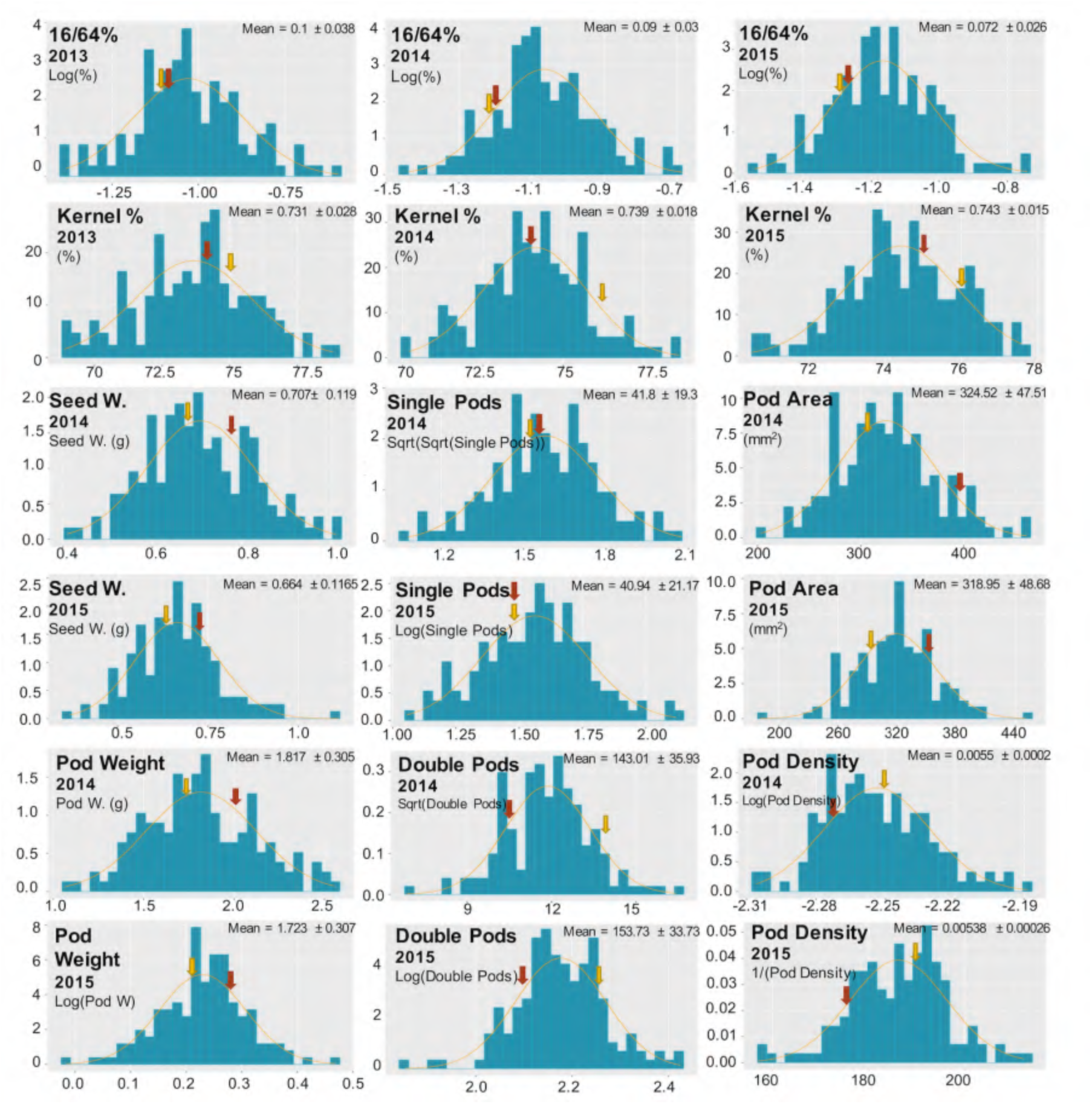
Phenotypic distribution for all traits in three consecutive years. Y-axis corresponds to density, and X-axis corresponds to the original metric or the normal-transformed trait value as indicated in the left corner or each plot, based on the average of the three replicates per year. Log, logarithm; Sqrt, square root; 1/, reciprocal. Arrows indicate the phenotypic values for NC 3033 (red) and Tifrunner (yellow). A normal distribution curve is shown in orange. The mean and SD values are based on the raw data according to Table 1.

**Table 1.** Summary statistics for seed and pod traits in parents and the RILs based on raw data.

|  |  | Parents |  | RILs |  |  |
| --- | --- | --- | --- | --- | --- | --- |
| | Variable | Tifrunner | NC 3033 | Mean ± SD | Minimum | Maximum |
| 2013 | 16/64P (%) | 8.007 | 8.093 | 10 ± 3.8 | 4.133 | 25.43 |
| | KP (%) | 75 | 74 | 73.1 ± 2.8 | 61.315 | 78.837 |
| 2014 | 16/64P (%) | 6.87 | 6.94 | 9.0 ± 3.0 | 3.644 | 20.564 |
| | KP (%) | 76 | 74 | 73.9 ± 1.8 | 66.223 | 78.599 |
| | PW (g) | 1.68 | 2.05 | 1.817 ± 0.305 | 1.067 | 2.5516 |
| | SdW (g)* | 0.67 | 0.79 | 0.707 ± 0.119 | 0.4107 | 1.00975 |
| | SP (count) | 30.00 | 32.33 | 41.8 ± 19.3 | 12 | 118.67 |
| | DP (count) | 164.33 | 103.67 | 143.01 ± 35.93 | 46.67 | 269.67 |
| | PA (mm<sup>2</sup>) | 301.22 | 389.55 | 324.52 ± 47.51 | 204.02 | 460.59 |
| | PD (g/mm<sup>2</sup>) | 0.0056 | 0.0053 | 0.0055 ± 0.0002 | 0.004946 | 0.006477 |
| 2015 | 16/64P (%) | 5.01 | 5.04 | 7.2 ± 2.6 | 2.863 | 18.139 |
| | KP (%) | 76 | 75 | 0.743 ± 0.015 | 69..268 | 77.567 |
| | PW (g) | 1.62 | 1.90 | 1.723 ± 0.307 | 0.9631 | 2.9426 |
| | SdW (g)* | 0.64 | 0.75 | 0.664 ± 0.1165 | 0.3681 | 1.11175 |
| | SP | 23.33 | 23.33 | 40.94 ± 21.17 | 11.33 | 132 |
| | DP | 178.67 | 126.67 | 153.73 ± 33.73 | 71.33 | 265.67 |
| | PA (mm<sup>2</sup>) | 285.81 | 360.12 | 318.95 ± 48.68 | 185.57 | 520.9 |
| | PD (g/mm<sup>2</sup>) | 0.00568 | 0.00527 | 0.00538 ± 0.00029 | 0.004714 | 0.006371 |

16/64P, 16/64 percentage as seed size index; KP, kernel percentage; PW, pod weight; SdW, seed weight; SP, single-kernel pods; DP, double-kernel pods; PA, pod area; PD, pod density.
\*SdW is reported as individual seed by dividing the original value from the weight of the two seeds contained in a pod.

The two parents contrasted for traits, Tifrunner was higher for KP, 16/64P, DP and PD, whereas, NC 3033 was higher for SdW, PW and PA. The population exhibited variation for all traits (Fig. 3), suitable for statistical and QTL analysis. Based on the analysis of variance and the boxplots for all the RILs by blocks (replicates) in all the years, we found block effects (Fig. 2), especially for 16/64P and KP in 2014 and 2015, SP and PA in 2014, and SdW and PD in 2015. Analysis of variance of all traits revealed significant differences between RILs and between years except for SP and PA, and the year x RIL interaction except for SP where there was no significant difference (Table 2). The broad sense heritability ranged from 61.3% to 80.3% for most of the traits, except for SP with a value of 40.4%, indicating a genetic component underlying these traits in this population (Table 2).

**Table 2.** Analysis of variance and heritability for seed and pod traits for the RIL population across three years.

| Trait | Variables | df | Mean Square | F-value | P-value | h<sup>2</sup> |
| --- | --- | --- | --- | --- | --- | --- |
| **16/64P** | Year | 2 | 1.773000 | 66.732 | <0.001 | 74.4% |
|  | RIL | 158 | 0.146410 | 22.969 | <0.001 |  |
|  | RIL x Year | 283 | 0.010780 | 1.692 | <0.001 |  |
|  | Error | 771 | 0.006370 |  |  |  |
| **KP** | Year | 2 | 0.006400 | 13.101 | <0.001 | 68.7% |
|  | RIL | 158 | 0.002480 | 20.106 | <0.001 |  |
|  | RIL x Year | 283 | 0.000295 | 2.390 | <0.001 |  |
|  | Error | 772 | 0.000123 |  |  |  |
| **PW** | Year | 1 | 0.133000 | 20.177 | <0.001 | 79.3% |
|  | RIL | 154 | 0.029922 | 19.764 | <0.001 |  |
|  | RIL x Year | 151 | 0.002186 | 1.444 | <0.005 |  |
|  | Error | 575 | 0.001514 |  |  |  |
| **SdW** | Year | 1 | 0.172000 | 26.291 | <0.001 | 80.3% |
|  | RIL | 154 | 0.030020 | 21.008 | <0.001 |  |
|  | RIL x Year | 151 | 0.002080 | 1.456 | <0.05 |  |
|  | Error | 575 | 0.001429 |  |  |  |
| **SP** | Year | 1 | 0.021000 | 0.212 | NS | 61.3% |
|  | RIL | 154 | 0.330870 | 7.424 | <0.001 |  |
|  | RIL x Year | 150 | 0.047960 | 1.076 | NS |  |
|  | Error | 561 | 0.044570 |  |  |  |
| **DP** | Year | 1 | 0.000122 | 11.281 | <0.001 | 40.4% |
|  | RIL | 154 | 0.000025 | 3.429 | <0.001 |  |
|  | RIL x Year | 151 | 0.000010 | 1.347 | <0.01 |  |
|  | Error | 576 | 0.000007 |  |  |  |
| **PA** | Year | 1 | 0.014504 | 3.058 | NS | 79.6% |
|  | RIL | 154 | 0.021535 | 19.677 | <0.001 |  |
|  | RIL x Year | 151 | 0.001464 | 1.337 | <0.01 |  |
|  | Error | 575 | 0.001094 |  |  |  |
| **PD** | Year | 1 | 0.000009 | 73.824 | <0.001 | 67.5% |
|  | RIL | 154 | 0.000000 | 11.275 | <0.001 |  |
|  | RIL x Year | 151 | 0.000000 | 1.751 | <0.001 |  |
|  | Error | 571 | 0.000000 |  |  |  |

h<sup>2</sup>, broad sense heritability.
NS indicates non-significance at P-value < 0.05.

Pearson correlations between traits (Fig. 4) were, as expected, mostly correlated, particularly for traits such as SdW, PW, PA and PD. Some traits had negative correlations such as 16/64P with KP, SdW, PW, PA and PD, also expected. In addition, SP was negatively correlated with KP, SdW, PW and PA, and DP negatively correlated with PW, SdW, PA, KP and PD in 2014-2015. There was some year to year variation as in 2013 KP was not correlated with other traits such as PW, DP, PA and PD in 2014-2015.

**Figure 4.**
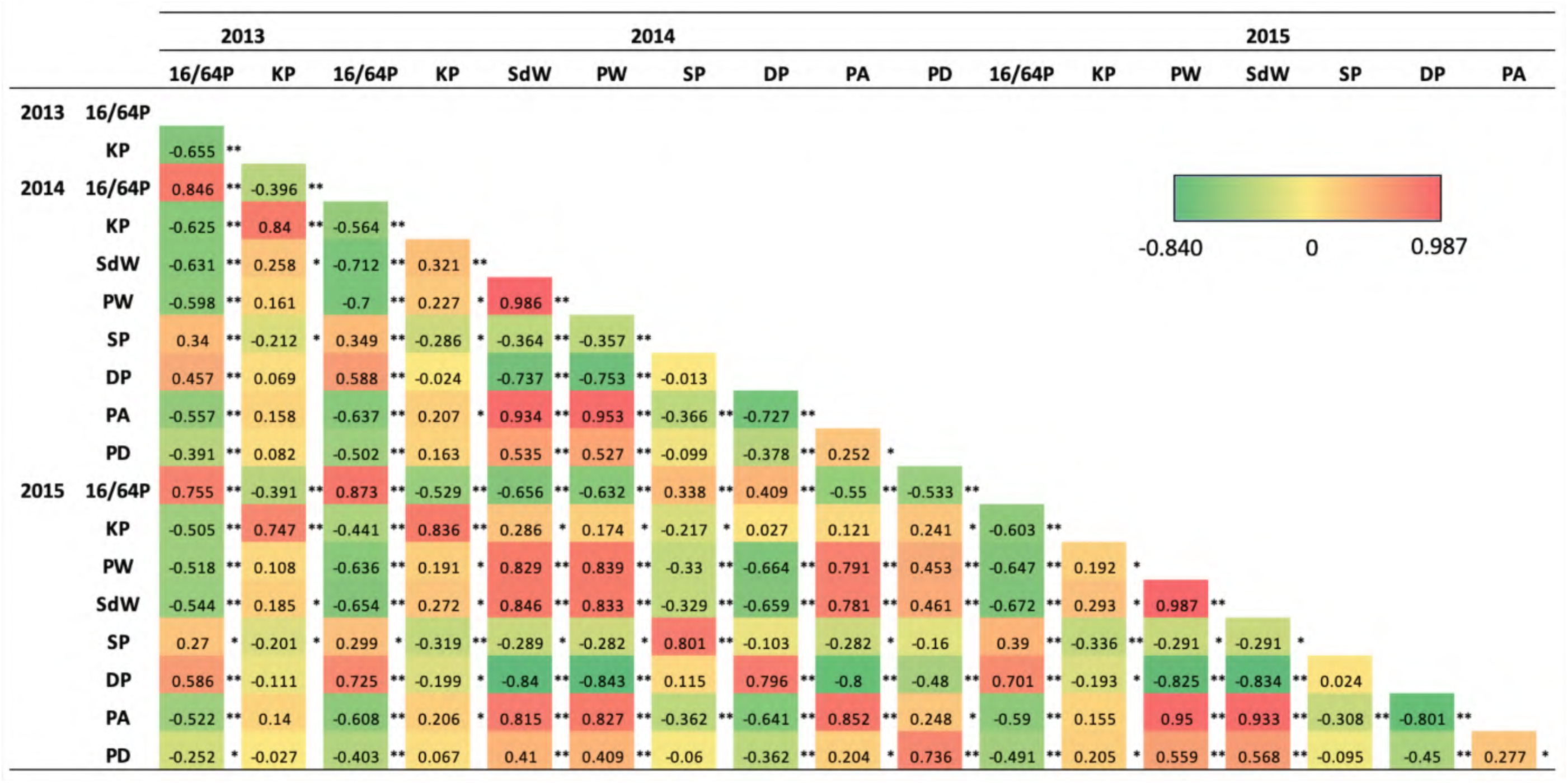
Pearson correlations for the seed and pod traits evaluated over three years. Red for the highest value and dark green for lowest value on the heatmap scale. Significant correlations * P < 0.05 and ** P < 0.001. 16/64P, 16/64 percentage as seed size index; KP, kernel percentage; PW, pod weight; SdW, seed weight; SP, single-kernel pods; DP, double-kernel pods; PA, pod area; PD, pod density.

### Linkage map and comparison with physical map

Genotyping of Tifrunner x NC 3033 RILs resulted in 2,233 polymorphic SNPs. After filtering for missing data and heterozygous calls, 1,998 SNPs and 100 SSRs were retained and a genetic map was constructed using the 156 selected RILs. The total map size spanned 3382.0 cM containing 1524 markers (1451 SNPs and 73 SSRs) assigned to 29 linkage groups (Fig. 5 and Table 3); 10 were from the A genome, 13 from the B genome and 6 were A and B markers combined. The 29 linkage groups ranged in size from A04 covering 298.7 cM to A08_B08 with 4.5 cM total with an average number of loci per linkage group of 53 ranging up to 133 loci in A04. The average distance between neighboring markers was 2.7 cM, ranging from 1.0 cM in B06_2 to 6.2 cM in B03.

**Figure 5.**
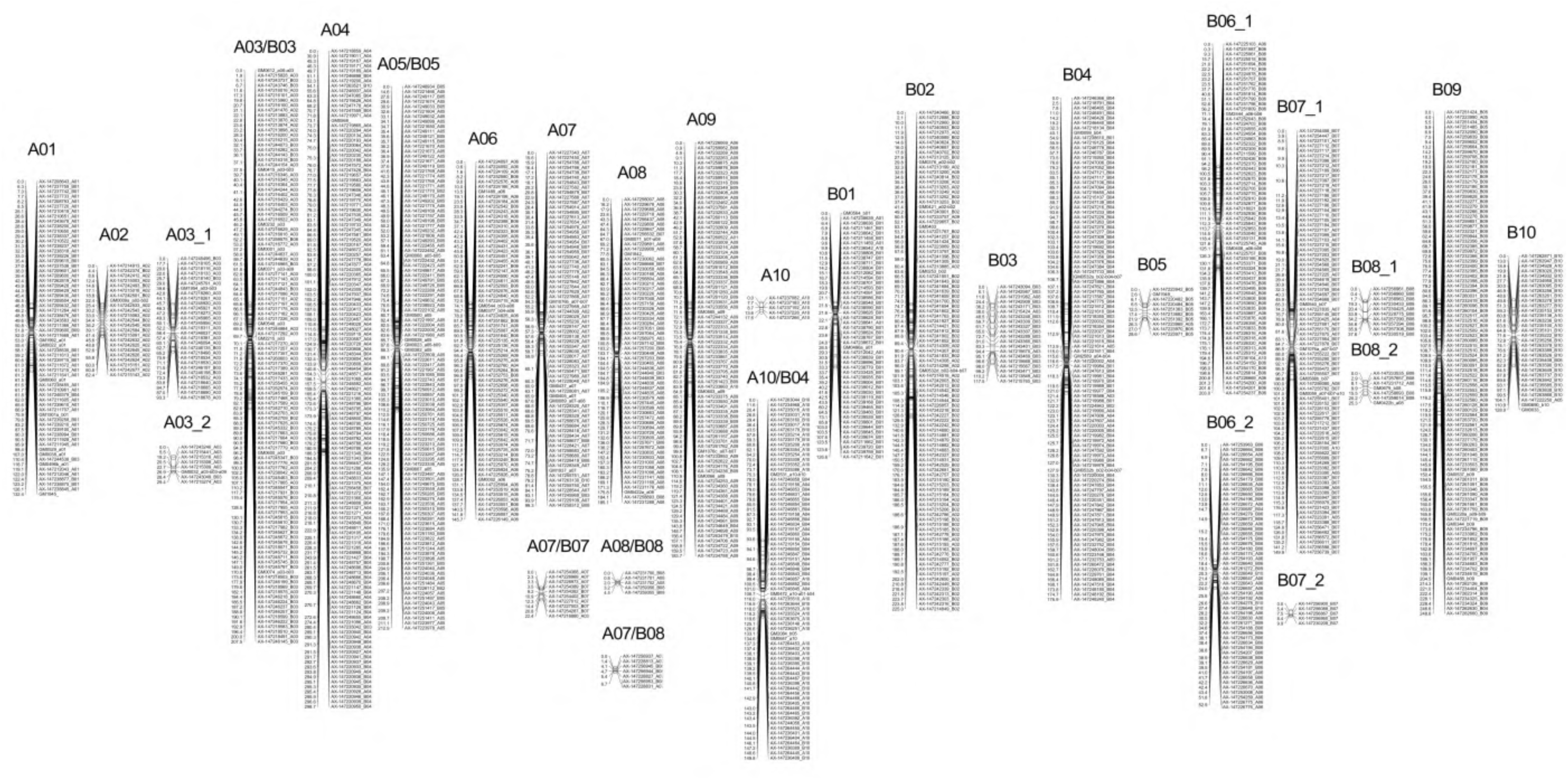
Genetic linkage map for the Tifrunner x NC 3033 RIL population. Distance in centimorgan (cM) is shown on left side of each group. The names of the SNPs are followed by the original chromosome number assigned when they were described. The name of the linkage groups was assigned based on the tetraploid reference genome.

**Table 3.**
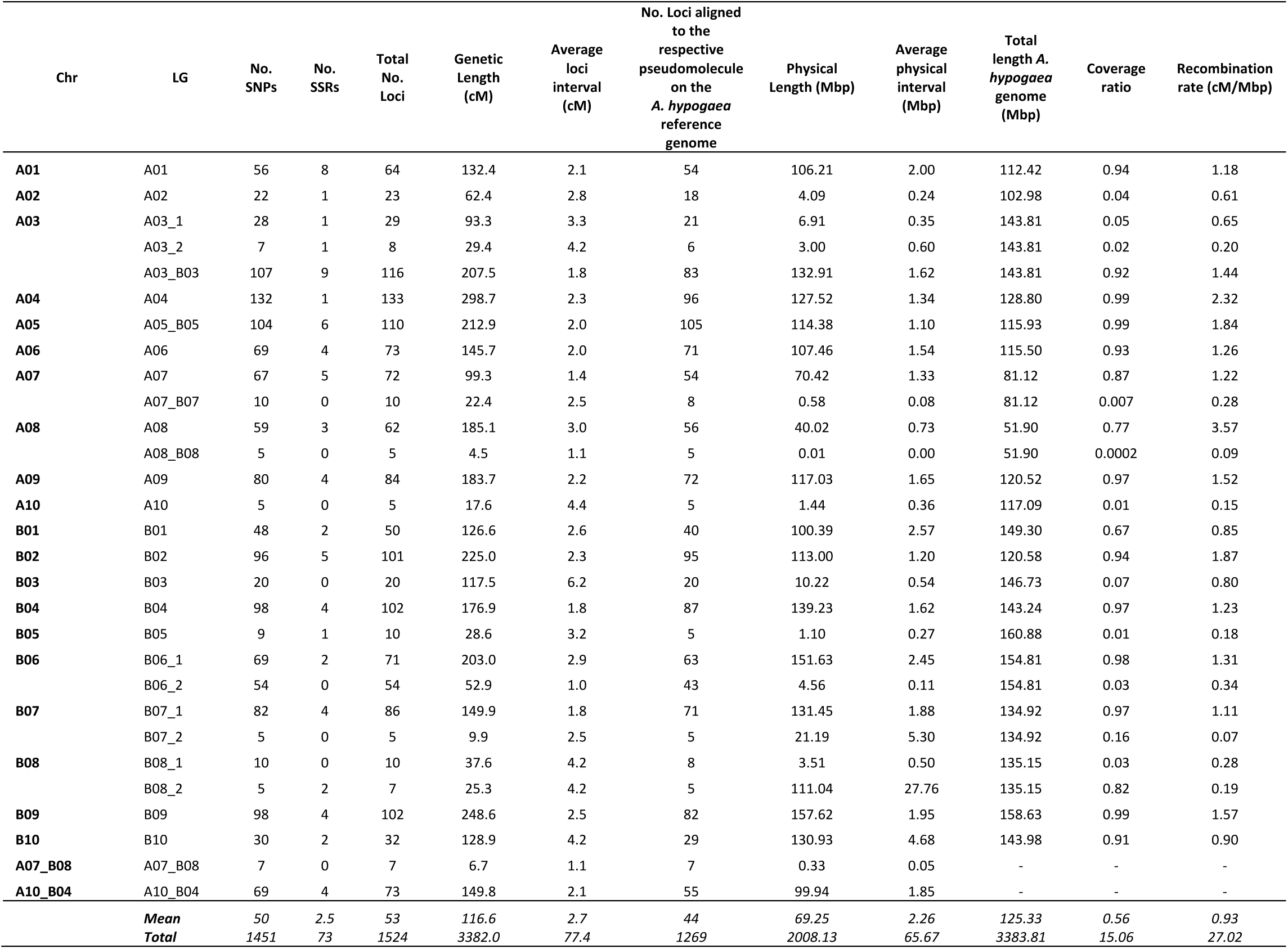
Genetic map description and comparison with physical distance based on the *A. hypogaea* reference genome.

| Chr | LG | No. SNPs | No. SSRs | Total No. Loci | Genetic Length (cM) | Average loci interval (cM) | No. Loci aligned to the respective pseudomolecule on the *A. hypogaea* reference genome | Physical Length (Mbp) | Average physical interval (Mbp) | Total length *A. hypogaea* genome (Mbp) | Coverage ratio | Recombination rate (cM/Mbp) |
| --- | --- | --- | --- | --- | --- | --- | --- | --- | --- | --- | --- | --- |
| A01 | A01 | 56 | 8 | 64 | 132.4 | 2.1 | 54 | 106.21 | 2.00 | 112.42 | 0.94 | 1.18 |
| A02 | A02 | 22 | 1 | 23 | 62.4 | 2.8 | 18 | 4.09 | 0.24 | 102.98 | 0.04 | 0.61 |
| A03 | A03_1 | 28 | 1 | 29 | 93.3 | 3.3 | 21 | 6.91 | 0.35 | 143.81 | 0.05 | 0.65 |
|  | A03_2 | 7 | 1 | 8 | 29.4 | 4.2 | 6 | 3.00 | 0.60 | 143.81 | 0.02 | 0.20 |
|  | A03_B03 | 107 | 9 | 116 | 207.5 | 1.8 | 83 | 132.91 | 1.62 | 143.81 | 0.92 | 1.44 |
| A04 | A04 | 132 | 1 | 133 | 298.7 | 2.3 | 96 | 127.52 | 1.34 | 128.80 | 0.99 | 2.32 |
| A05 | A05_B05 | 104 | 6 | 110 | 212.9 | 2.0 | 105 | 114.38 | 1.10 | 115.93 | 0.99 | 1.84 |
| A06 | A06 | 69 | 4 | 73 | 145.7 | 2.0 | 71 | 107.46 | 1.54 | 115.50 | 0.93 | 1.26 |
| A07 | A07 | 67 | 5 | 72 | 99.3 | 1.4 | 54 | 70.42 | 1.33 | 81.12 | 0.87 | 1.22 |
|  | A07_B07 | 10 | 0 | 10 | 22.4 | 2.5 | 8 | 0.58 | 0.08 | 81.12 | 0.007 | 0.28 |
| A08 | A08 | 59 | 3 | 62 | 185.1 | 3.0 | 56 | 40.02 | 0.73 | 51.90 | 0.77 | 3.57 |
|  | A08_B08 | 5 | 0 | 5 | 4.5 | 1.1 | 5 | 0.01 | 0.00 | 51.90 | 0.0002 | 0.09 |
| A09 | A09 | 80 | 4 | 84 | 183.7 | 2.2 | 72 | 117.03 | 1.65 | 120.52 | 0.97 | 1.52 |
| A10 | A10 | 5 | 0 | 5 | 17.6 | 4.4 | 5 | 1.44 | 0.36 | 117.09 | 0.01 | 0.15 |
| B01 | B01 | 48 | 2 | 50 | 126.6 | 2.6 | 40 | 100.39 | 2.57 | 149.30 | 0.67 | 0.85 |
| B02 | B02 | 96 | 5 | 101 | 225.0 | 2.3 | 95 | 113.00 | 1.20 | 120.58 | 0.94 | 1.87 |
| B03 | B03 | 20 | 0 | 20 | 117.5 | 6.2 | 20 | 10.22 | 0.54 | 146.73 | 0.07 | 0.80 |
| B04 | B04 | 98 | 4 | 102 | 176.9 | 1.8 | 87 | 139.23 | 1.62 | 143.24 | 0.97 | 1.23 |
| B05 | B05 | 9 | 1 | 10 | 28.6 | 3.2 | 5 | 1.10 | 0.27 | 160.88 | 0.01 | 0.18 |
| B06 | B06_1 | 69 | 2 | 71 | 203.0 | 2.9 | 63 | 151.63 | 2.45 | 154.81 | 0.98 | 1.31 |
|  | B06_2 | 54 | 0 | 54 | 52.9 | 1.0 | 43 | 4.56 | 0.11 | 154.81 | 0.03 | 0.34 |
| B07 | B07_1 | 82 | 4 | 86 | 149.9 | 1.8 | 71 | 131.45 | 1.88 | 134.92 | 0.97 | 1.11 |
|  | B07_2 | 5 | 0 | 5 | 9.9 | 2.5 | 5 | 21.19 | 5.30 | 134.92 | 0.16 | 0.07 |
| B08 | B08_1 | 10 | 0 | 10 | 37.6 | 4.2 | 8 | 3.51 | 0.50 | 135.15 | 0.03 | 0.28 |
|  | B08_2 | 5 | 2 | 7 | 25.3 | 4.2 | 5 | 111.04 | 27.76 | 135.15 | 0.82 | 0.19 |
| B09 | B09 | 98 | 4 | 102 | 248.6 | 2.5 | 82 | 157.62 | 1.95 | 158.63 | 0.99 | 1.57 |
| B10 | B10 | 30 | 2 | 32 | 128.9 | 4.2 | 29 | 130.93 | 4.68 | 143.98 | 0.91 | 0.90 |
| A07_B08 | A07_B08 | 7 | 0 | 7 | 6.7 | 1.1 | 7 | 0.33 | 0.05 | - | - | - |
| A10_B04 | A10_B04 | 69 | 4 | 73 | 149.8 | 2.1 | 55 | 99.94 | 1.85 | - | - | - |
| Mean |  | 50 | 2.5 | 53 | 116.6 | 2.7 | 44 | 69.25 | 2.26 | 125.33 | 0.56 | 0.93 |
| Total |  | 1451 | 73 | 1524 | 3382.0 | 77.4 | 1269 | 2008.13 | 65.67 | 3383.81 | 15.06 | 27.02 |

The names of the linkage groups were assigned based on the assignment of SNPs to the sequence-based pseudomolecules. If more than 51% of the markers were assigned to a specific chromosome it was given that name. In cases where the group contained ∼ 50% of loci from two chromosomes, the name included both chromosomes. Most linkage groups included markers from homoeologous chromosomes, however, two had markers from different chromosomes, A07_B08 and A10_B04 with 7 and 73 markers, respectively.

1,269 loci were successfully aligned to the *A. hypogaea* pseudomolecules spanning a total physical distance of 2008.13 Mbp and an average physical interval of 2.26 Mbp between loci (Table 3 and Fig. 6). The percentage of pseudomolecules covered by linkage maps varied, two groups covered more than 80% of the pseudomolecule, 12 groups more than 90% of which three were close to 100%, e.g. A04, A05_B05 and B09. The average recombination rate was 0.93 cM/Mbp and A08 had the maximum rate. A10, B05, B08_2 and A03_2 had the lowest recombination rates.

**Figure 6.**
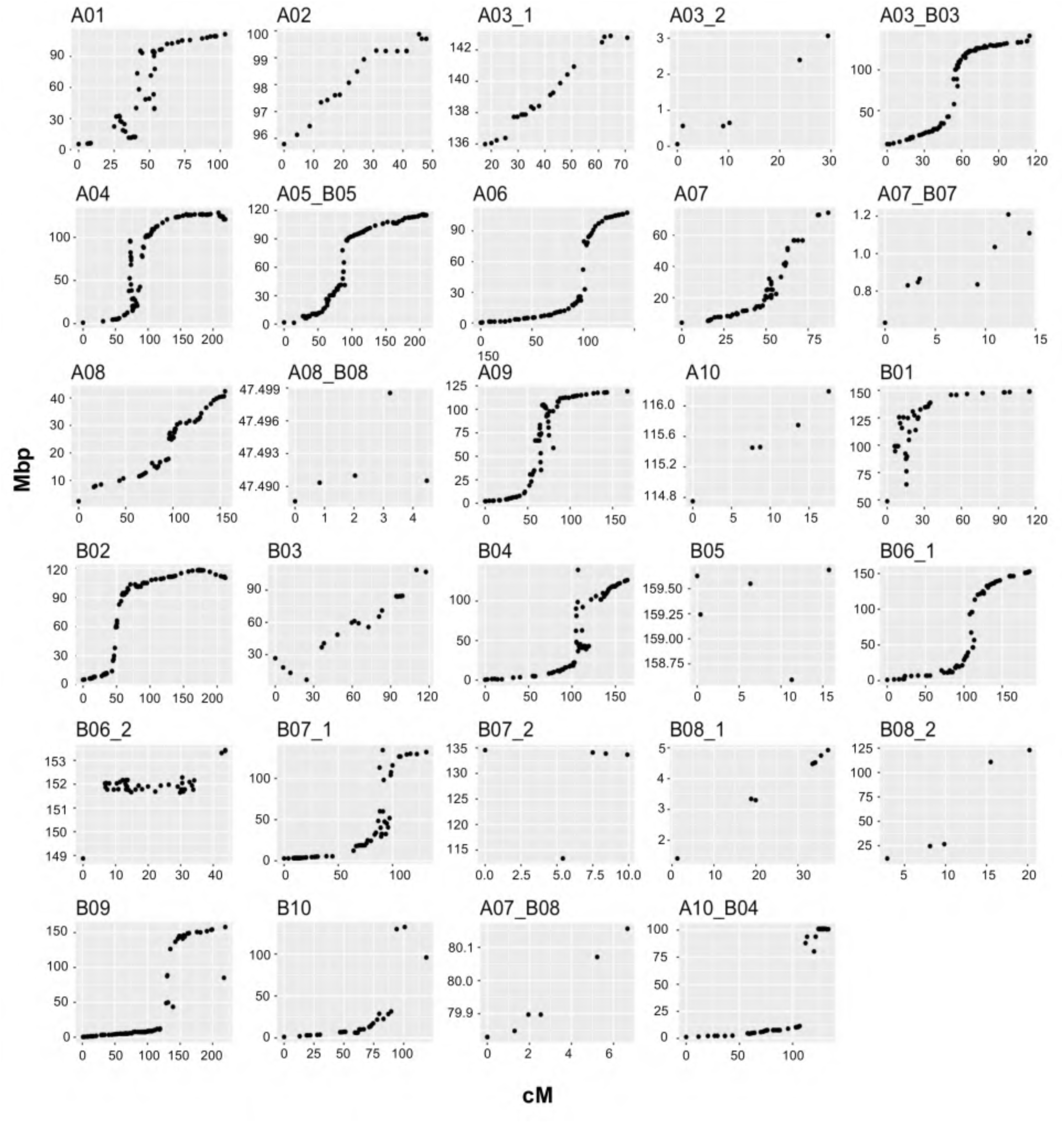
Genetic distance (cM) on x-axis vs physical distance in Mbp on y-axis for the Tifrunner x NC 3033 population based on the alignment of the SNP flanking sequences to the *A. hypogaea* reference genome cv. Tifrunner.

From the distribution of the loci along the chromosomes (Fig. 6) we observed higher marker saturation and increased recombination in the arms and lower marker saturation and recombination frequencies in the pericentromeric regions. Most of the linkage groups with good correspondence to a pseudomolecule were symmetrical, that is arms with dense markers and a pericentromeric region with few markers and reduced recombination. A few linkage groups exhibited rearrangements such as A01 and B03 where there is an apparent inversion on the top arm. Even though the marker density was low, there was a correspondence between loci from the group A07_B08 with the A07 pseudomolecule, as suggested previously (Bertioli *et al*. 2016).

### QTL identification

For seed and pod phenotypes, we identified 49 QTL on 14 linkage groups (Table 4 and Fig. 7). Most linkage groups had only one or two QTL, with a maximum of 14 QTL in A04, 11 QTL in A07_B07 and 10 QTL in B06_1. QTL were identified for all traits (16/64P, KP, PW, SdW, SP, DP, PA and PD) across all years, except for 16/64P in 2014 and 2015, and the QTL explained 5.3% to 31.4% of the phenotypic variation (Table 4). Eight QTL were major, explaining > 20% of the phenotypic variation, and 12 QTL had effects ranging between 10-20%. NC 3033 contributed most, 6 of 8, of the major QTL, all on B06_1, accounting for 24.4% - 31.4% of phenotypic variation. Tifrunner contributed two major QTL on B06_1 and A07_B07 corresponding to 28.4% and 29.2% of the phenotypic variation, respectively. Seven of the major QTL were associated with just two SNP markers, AX-147226319_A06 and AX-147226313_A06, that are 3.3 cM apart. These QTL were detected for four traits, PW, SdW, PA and DP, for years 2014 and 2015. The first three QTL were contributed by NC 3033 and had high positive correlations (Fig. 4), but were all negatively correlated with DP, contributed by Tifrunner. One QTL (*qDPA07_B07.2*) was located on A07_B07 (Table 4).

**Figure 7.**
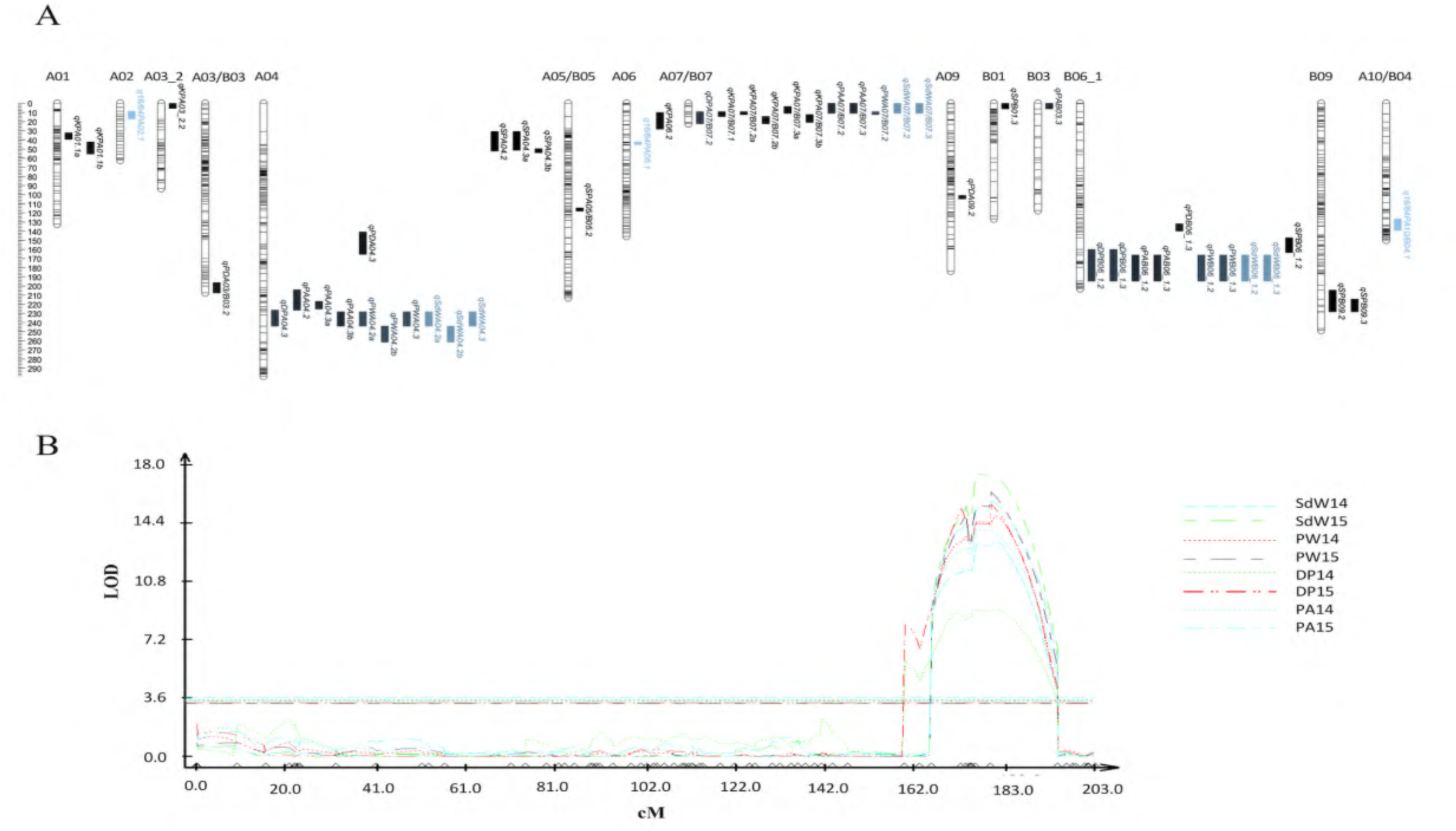
Overview of QTL identified for seed and pod traits on the Tifrunner x NC 3033 population. **A.** Linkage groups of the genetic map with QTL positions indicated. The QTL identified for all the traits are differentiated by color. **B.** Linkage group B06_1 indicating the QTL co-localizing on the bottom arm of the group. The y-axis represents the LOD score and the x-axis represents the distance (cM) of the linkage group and the markers mapped indicated by triangles on the bottom axis.

**Table 4.**
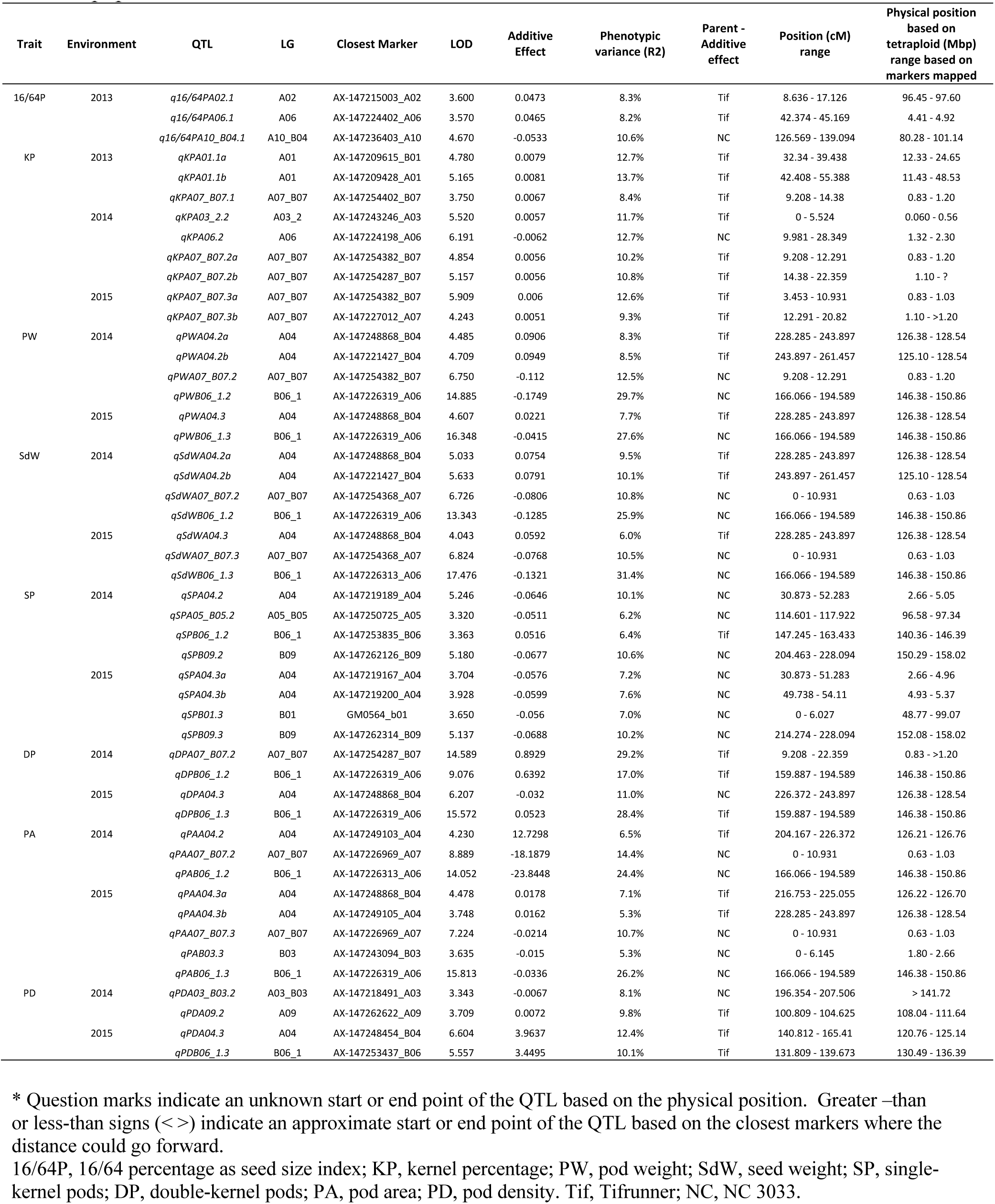
QTL information for seed and pod traits in peanut across the three years in the RIL population

KP had the most QTL, 9 over all three years, 8 were contributed by Tifrunner and one from NC 3033. NC 3033 contributed seven of nine SP QTL, three of them in A04. For PA, five of nine QTL were contributed by NC 3033. Seven QTL were identified for SdW with two major QTL on B06_1 provided by NC 3033 explaining 25.9% and 31.45% of the phenotypic variation. Six and four QTL were identified for PW and DP, respectively, on B06_1 with large effects (17.0% - 29.2%). For 16/64P, three QTL were found, two from Tifrunner on chromosomes A02 and A06, and one on A10_B04 from NC 3033. Finally, four QTL were found for PD, one from NC 3033 on A03_B03 and three from Tifrunner on A09, A04 and B06_1.

### Genomic positions and co-localization of QTL

The genetic positions of QTL in cM correspond to the end points where peaks exceeded statistical thresholds based on permutation tests. The approximate physical positions of the QTL were defined as the closest flanking genetic markers (Table 4). The average genetic distance spanned by the QTL was 15 cM corresponding to an average of 4.76 Mbp physical distance, though some ranged up to 50.3 Mbp. We observed that some QTL spanned similar genetic regions, in particular those on A04, A07_B07, B06_1 and B09 (Table 5).

**Table 5.**
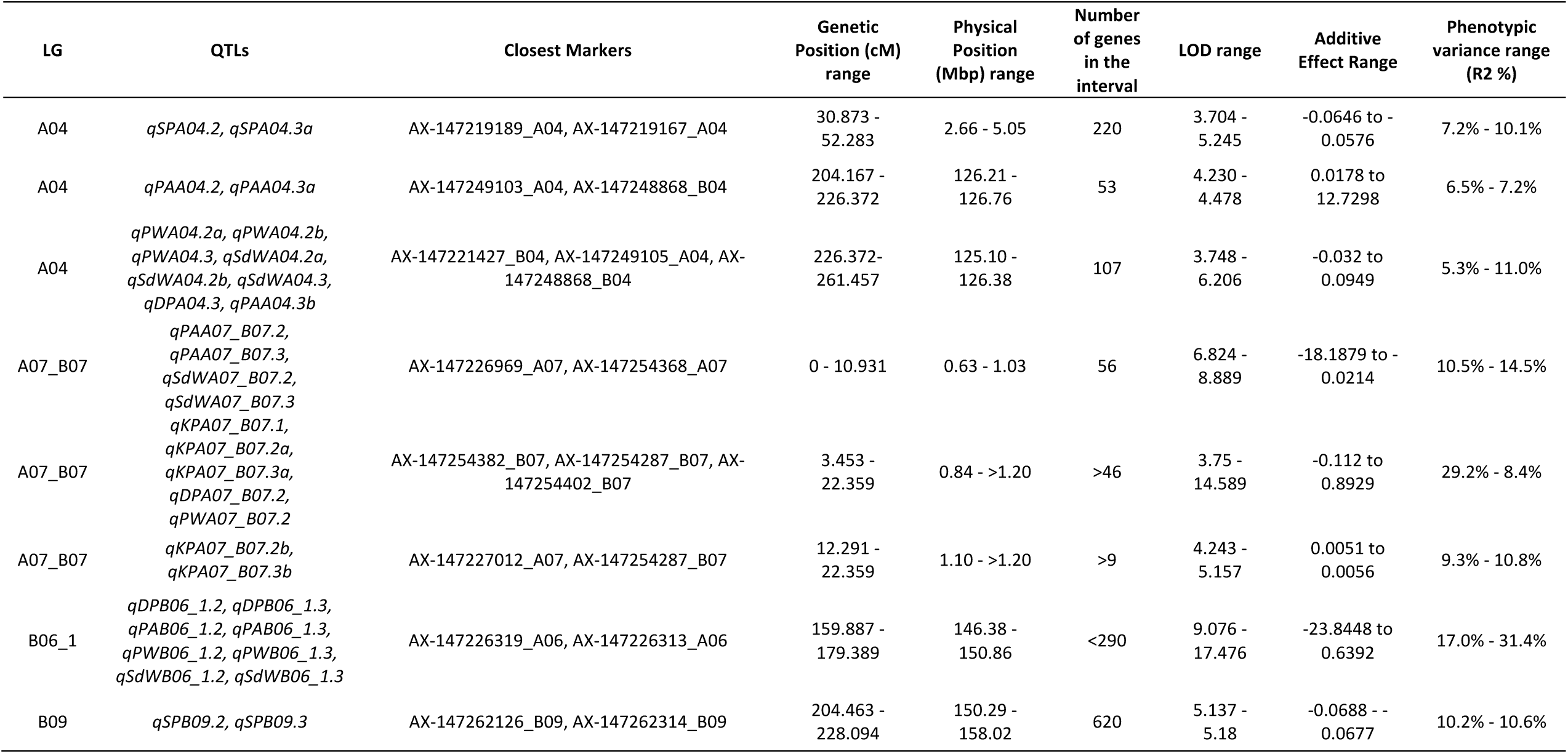
Co-localizing QTL including a range of genetic and physical positions and number of genes comprised on the respective regions.

We observed extensive clustering of QTL, as might be expected given the traits and correlations. On A04, three groups of QTL were co-localized, two of them overlapping between them. The first group included two QTL for SP, the second group two for PA, and the third group included 8 QTL: three for PW, three for SdW, and one each for DP and PA (Table 5). There are 220, 53, and 107 annotated genes within the physical regions spanned by the QTL, respectively. On A07_B07, another three QTL groups overlapped: the first group included two QTL each for PW and SdW; the second group included three QTL for KP and one for DP and PW, and the third group included two QTL for KP. There were several common markers in the QTL regions for groups two and three as these two groups overlapped by about 10 cM. The first group spanned 56 genes and the second and third more than 46 genes (Table 5). Other QTL clusters were observed, including those on linkage groups B06_1 and B09.

Co-localization of QTL and correlation of traits may be explained by pleiotropic effects for pod and seed phenotypes. There was, as expected, a high correlation in the behavior of the same traits across different years, confirmed by co-localization of QTL. Some QTL were both co-localized and highly correlated with other traits such as for PW, SdW and PA on A04, PA and SdW on A07_B07, and PA, PW and SdW on B06_1.

### Comparison with previously reported QTL

The physical locations of the QTL found in this research were compared with previous QTL studies for seed and pod traits by Chen et al. (2016a, 2017, 2019), Fonceka et al. (2012), Gomez Selvaraj et al. (2009), Wang et al. (2018a) and Luo et al. (2018) (Table 6 and S1). For 81 QTL from these seven studies, we were able to find either the marker sequences (Gomez Selvaraj *et al*. 2009; Fonceka *et al*. 2012; Chen *et al*. 2016a, 2017; Luo *et al*. 2018), or the sequence of the entire QTL from the two diploid progenitors (Wang *et al*. 2018a; Chen *et al*. 2019) and determined their positions by sequence alignment using BLAST to the reference genome.

**Table 6.**
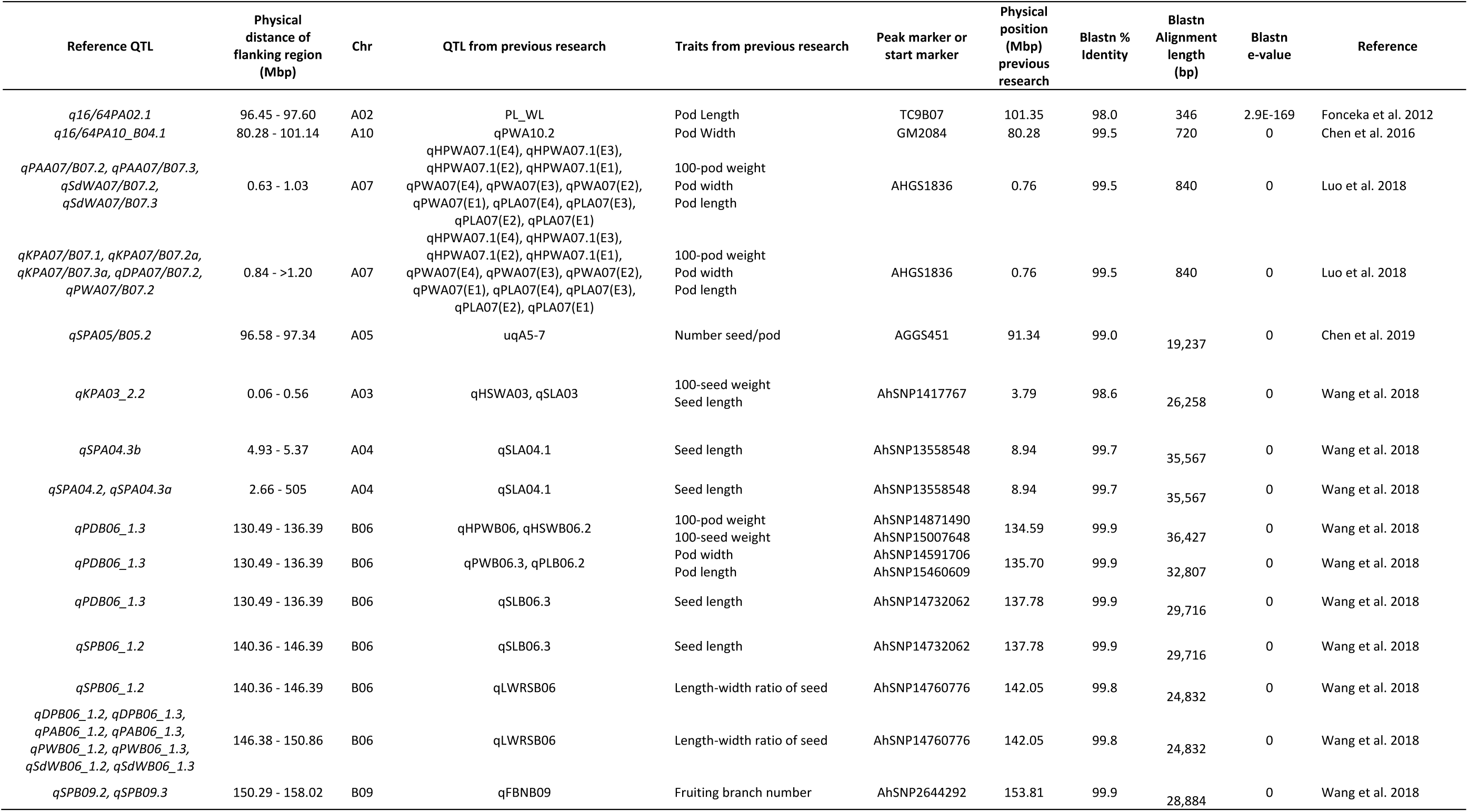
QTL found close or overlapping with QTL identified for seed and pod traits in previous studies. The comparison was made based on the physical location of the tetraploid species by sequence alignment using BLAST to the reference genome.

After the comparison with the QTL regions from previous studies, we found 11 QTL in close proximity (0.08 Mbp – 5.24 Mbp) on chromosomes A02, A03, A04, A05, A07, A09 and B06 and 6 QTL co-localizing in A07, A10, B06 and B10 (Table 6). No overlapping QTL were found for Selvaraj et al. (2009), but one from Fonceka et al. (2012), Chen et al. (2017), Chen et al. (2019), Luo et al (2018) and six from Wang et al. (2018) were found in close proximity to QTL from this study.

In comparison to Chen et al. (2016), one of our QTL co-localized with theirs at 80.28 Mbp of A10, which is close to the QTL flanking marker GM2084 (Genebank ID GO263349.1). In A07, the QTL cluster found by Luo et al. (2018) which included 12 QTL, co-localized with the QTL cluster found in this study around 0.63 – 1.03 Mbp linked to the marker AHGS1836 (Genebank ID_DH965050.1). Furthermore, four co-localizing regions were found after the comparison with the QTL discovered by Wang et al. (2018a), three of them at the bottom of the chromosome B06 (130.49 - 146.39 Mbp) and one in B09, including some QTL clusters (Table 6).

Due to the use of common markers, Chen et al. (2017) identified a group of unique QTL based on a comparison with previous studies (Gomez Selvaraj *et al*. 2009; Fonceka *et al*. 2012; Shirasawa *et al*. 2013; Pandey *et al*. 2014; Huang *et al*. 2015; Chen *et al*. 2016a, 2016b). After comparing the QTL from this research with the unique QTL reported by Chen et al. (2017), there was no evidence of overlapping QTL. However, there were some in close proximity (between 1Mbp - 4.8 Mbp) in the diploid genomes in chromosomes A02, B01 and B06.

## DISCUSSION

Approximately 3% of the markers on the SNP array were polymorphic in this population, reinforcing the observation that peanut has very low levels of sequence variation (Varshney *et al*. 2009; Hong *et al*. 2010; Chen *et al*. 2016a). As with other peanut studies, we had a high number of false positives in SNP calling due to the similarity between subgenomes (Clevenger *et al*. 2015, 2017; Clevenger and Ozias-Akins 2015). Thus, the low genetic polymorphism rate and genomic composition still thwart our ability to obtain high-quality, high-density maps obtained in other species. However, in comparison to previous studies, the number of markers in this map is quite high (Bertioli *et al*. 2014; Huang *et al*. 2016; Liang *et al*. 2017; Liu *et al*. 2019) and the distribution of the markers as compared to their physical positions in the tetraploid genome indicates reasonable coverage for QTL identification. Our map included 1,524 markers covering a map distance of 3,382 cM. The other five ‘high-density’ maps in peanut include 1,621 SNPs and 64 SSRs covering 1,446.7 cM (Zhou *et al*. 2014), 2,187 SNPs spanning 1,566.10 cM (Wang *et al*. 2018b), 3,630 SNPs covering 2,098.14 cM (Wang *et al*. 2018a), 3,693 markers in a consensus map spanning 2,651 cM (Shirasawa *et al*. 2013), and 8,869 SNPs (after whole genome population re-sequencing at 2x-5x coverage) with a map length of 3,120 cM (Agarwal *et al*. 2018).

Most of the SNPs were concordant with physical positions on the pseudomolecules, per their design (Pandey *et al*. 2017; Clevenger *et al*. 2017) and confirmed by sequence alignment after genetic mapping. For most linkage groups, it was possible to distinguish individual A and B genome chromosomes. However, there were six linkage groups (A03_B03, A05_B05, A07_B07, A08_B08, A07_B08 and A10_B04) where about 50% of the markers were assigned to the other sub-genome making it difficult to distinguish the A and B genome chromosomes. This is due to the high sequence similarity and collinearity between the A and B genomes and the low genetic diversity between them, due to a recent diversification of the two diploid progenitors (Bertioli *et al*. 2016).

Markers from A07 and B08 were in one linkage group corresponding to what Bertioli et al. (2016) described as a reciprocal translocation. A07 has a high repetitive content with only one euchromatic arm and A08 is a diminutive chromosome with high gene density (Bertioli *et al*. 2016). Thus, the physical composition of the chromosomes, and chromosome interchanges, may have played a role in the collapse of the genetic maps of these two groups as demonstrated by large syntenic blocks shared between A07 – B08 and B07 – A08.

Linkage maps were consistent with the new tetraploid sequences (Fig. 6) (Bertioli et al. 2019), which showed large inversions relative to the diploid genomes on A01, B01, B03 and B04 (Fig. 6). Bertioli et al. (2016) also found large inversions in both arms of chromosomes A01 and B01, and an apparent inversion in A05 as compared to the diploid reference genomes, also found by Wang et al. (2018a). These inversions were observed as an arc or a perpendicular line relative to the rest of the markers in a linkage group (e.g. A01 in Fig. 6), and in most cases, at the ends of the chromosome arms. These inversions likely drive DNA loss and/or gain through recombination-driven deletions that lead to DNA gain in non-recombinogenic regions (Bennetzen *et al*. 2005; Tian *et al*. 2009; Bertioli *et al*. 2016).

Although linkage groups did show some fragmentation compared to the chromosomal sequences, the markers were reasonably well distributed across the genome, based on genetic to physical distances and number per linkage group. Similar to other species, the pericentromeric regions were depauperate for markers and had low recombination rates (Jensen-Seaman *et al*. 2004; Sharma *et al*. 2013).

All the selected phenotypic traits demonstrated transgressive segregation, with some RILs showing extreme phenotypes and exceeding the performance of the parents, such as RILs PR F6:7_600, PR F6:7_620, PR F6:7_62, etc. (Table S2). Furthermore, the high broad sense heritability for all traits except DP indicated a major genetic component. Based on these observations, we inferred that this population was suitable for genetically dissecting seed and pod traits as a prelude to contributing to yield improvement.

In contrast to previous studies (Table S1), we used PD as a measurement for seed and pod filling and measured PA and PD based on methods described in Wu et al. (2015) in order to identify loci associated with these traits and to find correlations with traits measured in previous work. PD and PA had relatively low positive correlations demonstrating that large pods are not always associated with either larger seeds or higher yields. These results were expected as NC 3033 has larger pods than Tifrunner but has incomplete pod fill.

It was previously observed that large pods may be correlated with thick pericarp in peanut which complicates selection for large pods with large and dense seeds (Hammons 1973; de Godoy and Norden 1981; Venuprasad *et al*. 2011; Wu *et al*. 2015), and it was noted that the thickness of pods is highly correlated with pod maturity (Williams *et al*. 1987). This supports our finding of QTL co-localized on A07 for KP with previously mapped percentage of pod maturity (Fonceka *et al*. 2012). This demonstrates that maturity can be indirectly measured and that our population is likely segregating for maturity, since both parents of the population have different maturity ranges, Tifrunner being a late maturity peanut with ∼150 days after planting (Holbrook and Culbreath 2007) and NC 3033 with an earlier maturity of ∼135 days after planting (Beute *et al*. 1976; Korani *et al*. 2018). At the time of harvest, when seed and pod filling is complete and the seeds have accumulated storage products, the seed density is higher than in immature seeds (Williams *et al*. 1987; Sanders 1989; Rucker *et al*. 1994b). This is supported by high positive correlations of PD and PA with SdW and PW, demonstrating that it is possible to have larger pods and larger seeds. These results are also supported by Rucker et al. (Rucker *et al*. 1994a) showing that pods with mature kernels have significantly greater density. Although the population was segregating for duration of maturity, pod maturity was measured by the inner pericarp color to select samples for PW and SdW to calculate PA and PD and also to contrast the values with KP. In addition, we assumed there were no confounding effects with KP, since the correlations between KP vs SdW, PW, PA and PD were very low, and we could see an indirect measure of maturity from these traits.

On the other hand, PD and PA were negatively correlated with 16/64P, SP and DP, indicating that the larger pods with higher density had a smaller percentage of seeds passing through the screen. Tifrunner is a large-seeded runner type and NC 3033 a small-seeded Virginia type (Fig. 1). Regarding the negative correlation of PD and PA with SP and DP, this indicates that greater pod area and density are associated with lower pod count per standard sample weight, regardless of number of seeds in the pods. This corresponds to the co-localized QTL found for seed and pod weight vs single and double pods (Table 4, Fig. 4a).

This observation contrasts with work in Arabidopsis, however, where Gnan et al. (2014) found that seed number evolved independently from seed size due to a non-overlapping QTL found in a multiparental population, although natural variation is observed within the species. There are other studies corroborating the trade-off between seed size and seed number in crops when there are sufficient resources available at the time of seed set (Gambín and Borrás 2009). Furthermore, a correlation between seed number and the duration of seed filling period was observed (Kantolic and Slafer 2007) concordant with our findings that the population is likely segregating for maturity. Even though Tifrunner and NC 3033 are both characterized by double kernels, the population segregated for the number of seeds per pod with both single and double-kernel at a ratio of 1:4 single to double-kernel. This may also be explained, in part, by segregation for maturity in the population, related to the pod and seed filling period (Clarke 1979; Rucker *et al*. 1994b, 1994a; Kantolic and Slafer 2007; Gambín and Borrás 2009).

Regarding the distribution of QTL, Fonceka et al. (2012) identified 15 QTL on LG A07 and 17 QTL on B02 and B06, all for yield, seed and pod traits, with large phenotypic effects ranging from 8.7% to 26%, similar to this study. Wang et al. (2018a) found most of the QTL related to yield traits at the ends of B06 and B07 with phenotypic variation ranging from 4.30 - 18.99%, with six co-localized QTL in close proximity with QTL found in this study on B06 (Table 6).

Consequently, QTL related to seed and pod size and weight were concentrated on three linkage groups. This follows previous work suggesting that alleles from QTL for seed and pod size are clustered in A07, B02 and B06 due to domestication (Fonceka *et al*. 2012). Also, the seven QTL found in B06 confirmed previous studies, mainly the QTL found by Wang *et al*. (2018), which found pleiotropic QTL at the end of the B06 chromosome and found candidate genes associated with yield traits, some of them related to embryo development. These findings demonstrate the consistency of QTL across different genetic backgrounds and the potential for marker assisted selection of desirable seed and pod traits.

Of the 49 QTL identified, 33 co-localized with either the same trait in another year and/or with other traits in the same or different years. The regularity of the QTL discovered in the same linkage group locations across the years, the co-localization with previous studies, and the high phenotypic variance (Table 4, Table 5, Table 6) indicates the reliability of these QTL. Although the regions covered by the QTL are still large in physical distance, we were able to better elucidate the location of these QTL, including annotated genes in these regions that can be used to develop additional markers. Others have observed correlations between QTL regions with differentially expressed candidate genes, and it has been suggested that overlapping QTL might share common biochemical pathways (Schweizer and Stein 2011; Kocmarek *et al*. 2015); indeed many of the QTL in this study were correlated. Only a few QTL did not co-localize with others, even ones with high correlations, such as 16/64P and PD with r > 0.7.

In summary, we found new seed and pod QTL and validated QTL found in other populations. This provides additional tools for marker-assisted breeding to advance peanut improvement and for eventual molecular characterization of these economically important traits. Additional mapping is needed to further delineate the candidate genomic regions and find the genes causal to the phenotypic variation, and to pyramid the genes/QTL for superior genotypes. Marker assisted selection is in progress in peanut, currently used for only a few traits (Ozias-Akins *et al*. 2017); however, these QTL can expand the molecular breeding toolbox for peanut in order to improve the yield and quality of the peanut crop. To that end, marker-trait associations need to be further refined and validated in other breeding populations.

## Author contributions

Conceived and designed the experiments: POA, CCH, RH, SAJ. Population design: CCH, TGI. Performed the experiments: CC, YC, CCH. X-Ray measurements: CB, ML. Data analysis: CC, YC, DB. Writing/editing: CC, SAJ, POA, CCH.

## Acknowledgements

This work was supported by funding from the US-Israel Binational Agricultural Research and Development Fund (BARD IS-4540-12 to POA, RH and SAJ). We thank Jenny Leverett, Eric Antepenko, Caitlynn Schneider, Emma Matthews, James Watkins, Sirjan Sapkota, and Katherine Willard for their help with seed and pod phenotyping. We thank Shannon Atkinson, Jason Golden, Betty Tyler for the field work and X-Ray measurements in Tifton. We also thank Stephanie Botton and Kathleen Marasigan for their help with DNA extractions. We are additionally grateful to Jason Wallace and Chunming Xu for advice on statistical analysis and Soraya Bertioli, Chung-Jui Tsai and Ali Moussaoui for reviewing an early version of the manuscript.

